# Copy number evolution in simple and complex tandem repeats across the C57BL/6 and C57BL/10 inbred mouse lines

**DOI:** 10.1101/2021.03.01.433487

**Authors:** Jullien M. Flynn, Emily J. Brown, Andrew G. Clark

## Abstract

Simple sequence tandem repeats are among the most rapidly evolving compartments of the genome. Some repeat expansions are associated with mammalian disease or meiotic segregation distortion, yet the rates of copy number change across generations are not well known. Here, we use 14 distinct sub-lineages of the C57BL/6 and C57BL/10 inbred mouse strains, which have been evolving independently over about 300 generations, to estimate the rates of copy number changes in genome-wide tandem repeats. Rates of change varied across simple repeats and across lines. Notably, CAG, whose expansions in coding regions are associated with many neurological and other genetic disorders, was highly stable in copy number, likely indicating purifying selection. Rates of change were generally positively correlated with copy number, but the direction and magnitude of changes varied across lines. Some mouse lines experienced consistent losses or gains across most genome-wide simple repeats, but this did not correlate with copy number changes in complex repeats. Rates of copy number change were similar between simple repeats and the much more abundant complex repeats once they were normalized by copy number. Finally, the Y-specific centromeric repeat had a 4-fold higher rate of change than the homologous centromeric repeat on other chromosomes. Structural differences in satellite complexity, or restriction to the Y chromosome and the elevated mutation rate of the male germline, may explain the higher rate of change. Overall, our work underscores the mutational fluidity of long tandem arrays of repeats, and the correlations and constraints between genome-wide tandem repeats which suggest that turnover is not a completely neutral process.

## Introduction

Tandem arrays of repetitive DNA sequences are emerging as interesting and important parts of genomes (Lower *et al*. 2018). Like all other genomic compartments, tandem repeat sequences are subject to mutations, including single nucleotide mutations and copy number changes. The latter are often caused by replication slippage, unequal exchange, and rolling circle amplification (Smith 1976; Charlesworth *et al*. 1994; Cohen *et al*. 2003). Tandem repeats evolve at high rates, as demonstrated by the rapid turnover of repeats and abundance of differences between species (Wei *et al*. 2018; Cechova *et al*. 2019). Repeat copy number expansions, especially glutamine (CAG) arrays in coding regions, are involved in many human genetic disorders (reviewed in (Lieberman *et al*. 2019)). Even outside of coding regions, tandem repeat copy number changes could alter the phenotype, for example by acting as a sink for heterochromatin binding proteins and thereby influencing chromatin states and gene expression locally or genome-wide (Weiler and Wakimoto 1995; Lemos *et al*. 2010). Therefore the rate at which tandem repeats increase or decrease their copy number has relevance to genome biology and disease.

Tandem repeats can be broadly classified by the length of each repeat unit, which can range from a single nucleotide to hundreds or thousands of base pairs. In mammals, arrays of shorter repeat units (=<20 bp, simple repeats or kmers) tend to be located in both heterochromatic and euchromatic regions of mammalian genomes, whereas longer repeat units, generally >100 bp (also called complex repeats) make up the pericentromeric and centromeric regions. Tandem repeats are called satellite DNA when they form very long (at least several kilobase) arrays, and this is the case for the pericentromeric tandem repeats in mammals. In mouse, the total length of pericentromeric satellite arrays have been shown to influence the chromosome’s fate in meiosis, with longer arrays showing a transmission advantage compared to a homologous chromosome with shorter arrays (Iwata-Otsubo *et al*. 2017). In humans, there is also polymorphism in centromeric satellite copy number, although the repercussions on meiosis, DNA fragility, and transmission are not well understood (Black and Giunta 2018).

A few studies in Daphnia and Chlamydomonas have used mutation accumulation (MA) lines to study simple repeat mutation rates (Flynn *et al*. 2017, 2018; Ho *et al*. 2019). Ideally MA experiments employ large numbers of lines for many generations, with the lines maintained in a way that minimizes natural selection. Full ascertainment of mutations by whole-genome sequencing then provides an opportunity to estimate mutation rates, including rates of gains and losses in repeat copy number. These studies are much more cumbersome with mammals, which have longer generation times and are more labor-intensive and expensive to maintain.

The laboratory mouse, *Mus musculus,* is the most common mammalian model for genetic studies. The mouse genome is similar in size to the human genome and has an overall similar repeat structure (Consortium and Mouse Genome Sequencing Consortium 2002; Komissarov *et al*. 2011). The C57BL/6 (“Black 6”) mouse is the most commonly used inbred mouse strain used in biomedical research (The Jackson Laboratories). This strain and a related C57BL/10 (“Black 10”) line were created a century ago by C.C. Little (Festing 1979; Reeve 1989). Since then, different isolated subpopulations have been propagated in several different laboratories sequentially (Bailey 1978). Inbred mouse strains are maintained in a similar bottlenecking manner that maintains a low effective population size, thereby reducing selection and allowing mutations (except those causing sterility, lethality, or visible phenotypic defects) to accumulate. Each subpopulation can be treated as an independent MA experiment by combining the analysis of simple repeat content with a phylogenetic view of the inbred lines. In this way, rates of copy number change can be calculated based on reconstructing the ancestral state at each node and tallying the sequence differences between the nearest node and each branch tip.

In addition to copy number changes of individual repeats, we can study correlations in changes between repeats, which may indicate constraints on tandem repeat evolution. Although a traditional 20-generation MA study has been completed for mice which quantified the single nucleotide mutation rate (Uchimura *et al*. 2015), we made use of an inadvertent mutation-accumulation experiment in which strains of inbred mice diverged for many more generations and with a more complex phylogenetic structure in order to have greater ability to discover constraints among repeats. Previous work has shown evidence for trade-offs in copy number between repeats, indicated by negative correlations in copy number changes. These trade-offs are hypothesized to be mediated by maximum limits for repeat arrays (before inducing a negative phenotype) or co-localization of repeats adjacent to a functional sequence (Stephan and Cho 1994; Flynn *et al*. 2017). It is also possible for lines to greatly diverge in the magnitude and/or direction of copy number changes (Flynn *et al*. 2018). This is akin to “hypermutator lines” which have evolved a higher than ancestral mutation rate, typically in the context of single nucleotide mutations, which have been studied more widely. Finally, the dominant mechanisms (e.g. unequal exchange, replication slippage) that result in copy number changes for complex satellites may be different than that of simple tandem repeats. Since mice have abundant simple repeats and abundant well-characterized complex satellites (Wong and Rattner 1988; Vissel and Choo 1989; Komissarov *et al*. 2011), they provide a model system in which we can ask if rates of copy number change are similar or even correlated in both types of repetitive sequence.

Using genome mapping-based approaches to ascertain variation in simple repeat abundance among strains is challenging because most simple repeat-containing reads cannot be mapped to unique loci. Thus, we use a kmer identification and counting approach that does not require unique read-mapping to estimate simple repeat abundances in genomes. Although we cannot differentiate between loci of the same repeat with this approach, we can identify repeat-specific mutation rates in a less-biased manner. Here, we analyze simple and complex repeat copy number in unassembled short-read sequencing data from males of 14 B6/B10 mouse strains that have been propagated independently for 64-294 generations. We estimate copy number change rates of simple repeats as well as the major (pericentromeric), minor (centromeric), and Ymin (centromeric satellite specific to the Y) complex satellites.

## Materials and Methods

Relevant scripts used to analyze the data are available here: https://github.com/jmf422/BL6_mouse_satellites

### Mouse sequencing

Data used are from (Mortazavi *et al*. 2020). In brief, DNA from 1 male per strain was extracted from spleens. Sequencing libraries were prepared using a TruSeq DNA LT kit, as per the manufacturer’s instructions. The DNA was sequenced at Novogen at an average depth of 30X coverage on an Illumina HiSeq XTen (paired-end 150 bp reads).

### Simple repeat characterization

Adapters were trimmed with Trimmomatic. We ran k-Seek with standard parameters on the 14 mouse lines (Wei *et al*. 2014). In order to perform GC content and depth normalization of kmer abundances, reads were mapped to the mm10 genome with bowtie2. The GC correction pipeline from (Flynn *et al*. 2017) was run on each chromosome separately and then summed across chromosomes for normalization of kmer counts.

We also analyzed the current mouse genome assembly (version 39) for simple tandem repeats using Phobos (https://www.ruhr-uni-bochum.de/spezzoo/cm/cm_phobos.htm) to estimate the percent of genomic simple repeats we are able to capture with our approach. We limited the search to arrays that could theoretically be captured by k-Seek with short read sequencing (array length > 50 bp and unit size 1-20 bp).

### SNP analysis

In order to reduce the uncertainty in the time of common ancestry of the C57BL/6 and C57BL/10 strains, where historical records appear to be incomplete, we relied on divergence at single nucleotide sites, inferred from the genome sequences. Polymorphisms were called by GATK by following the recommended practices. We next removed indels from the VCF file to focus solely on single nucleotide polymorphisms (SNPs). We used the R package SNPRelate to process the vcf file. We calculated a distance matrix between samples using the SNP VCF file with the function snpgdsIBS, which calculates the proportion of sites that are identical-by-state in a pairwise manner among samples. Next we used snpgdsHCluster to hierarchically cluster the distance matrix followed by snpgdsCutTree to produce a tree. We did not use the SNP tree to reconstruct the relationships within the C57BL/6 and C57BL/10 clades since these are well documented by historical records, and the SNP-based phylogeny was consistent with these records (Mortazavi *et al*. 2020).

### Major and minor satellite analysis

We mapped the genomic reads to the major and minor satellites and estimated mean copy number based on the depth of reads mapping to the satellite and overall read depth. We mapped the reads to a multi-copy sequence of each satellite so that reads at the boundaries of tandem repeat copies would map. Since male mice were sequenced, we also analyzed the copy number of the Y-specific minor satellite (Ymin, (Pertile *et al*. 2009)). This repeat has a far different structure than the minor satellite, with complex higher order repeats (HORs) and each copy within the HOR being highly diverged. We therefore mapped reads to the HOR and calculated the rates on a per-HOR copy basis. We verified that no reads cross-mapped to both Ymin and the minor satellite. Scripts are available at: https://github.com/jmf422/BL6_mouse_satellites/tree/main/complex_satellites

### Phylogenetic modelling

We used the R package Phytools to construct a phylogeny of the 14 strains. First, we manually reproduced the phylogeny in .tre format. For all relationships within B6 and within B10, we used the historical records of known divergence times (Mortazavi *et al*. 2020). Since the history of the lines is recorded by the year of divergence, we used 4.33 generations per year to estimate the number of generations of each branch (Jackson Laboratories https://www.research.uci.edu/forms/docs/iacuc/JAX-breeding-strategies.pdf). We acknowledge that this generation time may be an overestimate for many of the laboratories, however this would not affect our conclusions because we used the same value for all lineages and we are mainly interested in the relative differences between lines. The genetic divergence between B6 and B10 clades was much greater than expected by their date of separation, suggesting that there was incomplete inbreeding at the time of separation or accidental introgression of haplotypes from another strain after their separation (Mortazavi *et al*. 2020). Based on our SNP analysis, the divergence level between B6 and B10 clades was proportional to 850 generations of separation on average across the genome (Figure S1), which is what we used to reproduce the tree. This generation number estimate near the base of the tree is not used for calculating the per generation rate of change of kmers but only for a more accurate estimate of kmer ancestral states at terminal nodes. We also have evidence from simple repeat abundances that the greatest magnitude of divergence/longest branches is between B6 and B10 (discussed below). We reconstructed the ancestral state, *A,* for the nearest node to the focal line, *m,* for each kmer, *k,* using the phytools fastAnc function, which calculates the estimated trait value at given nodes, assuming Brownian motion of trait values. We then subtracted the ancestral state copy number from the copy number for the focal kmer for the focal line. We then divide by the number of generations, *G,* between the ancestral node and focal line.

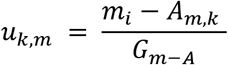

We use the terminology “copy number change rate” instead of “mutation rate” to reflect the possibility that the design of this study cannot accurately measure mutation rates. This is because the mouse phenotypes may be scrutinized more in inbred line maintenance than in a mutation accumulation study, thus purifying selection may play a role in maintaining copy numbers. Additionally, over the hundreds of generations since the ancestral node, the kmer abundances may have gone up and down in many mutational steps, and thus the net change in copy number is likely an underestimate of the true mutation rate. Since we are dealing with genome-wide simple repeats that are contained in an unknown number of arrays, and constraints may be acting in various ways, it is not possible to accurately model the mutation rate, so instead we only calculate the net copy number change rate.

Since copy number change rates tend to positively correlate with the copy number of the repeat (Flynn *et al*. 2017, 2018; Figure S3), we also calculated rates that are normalized by the copy number in the ancestral state. Copy number change rates are assigned a positive sign for gains in copy number, and a negative sign for a loss in copy number. We also calculated the absolute rates, which ignore direction and only reflect the absolute change in copy number. We show results for rates on a per-kmer basis and also a per-line basis.

It has been observed that kmer abundances sometimes do not mirror known phylogenetic relationships because of rapid changes in copy number possible in both positive and negative directions (Cechova et al. 2019). We asked whether for any kmers, the directional copy number changes propagated along the tree such that more closely related strains had more similar kmer abundances than distantly related strains (i.e. a phylogenetic signal). We used the package phylosig along with a manual approach (described below) to detect kmers with phylogenetic signal from the set of 427 total kmers. We verified each kmer with potential phylogenetic signal using the plot function in phytools. Our manual approach involved randomly assigning the ancestral node for the branch from which the terminal copy number was measured. If there was no phylogenetic signal, as was the case for most kmers, the mutation rate should average out to be very similar among kmers. Outlier kmers with highly negative or positive mutation rates were interpreted as likely having a phylogenetic signal. This manual approach allowed the identification of an additional kmer with a strong phylogenetic signal that the phylosig package did not detect. Scripts are available at: https://github.com/jmf422/BL6_mouse_satellites/tree/main/R_analysis.

## Results and Discussion

### Simple repeat characterization in the B6/B10 lines

We used k-Seek to characterize simple tandem repeats in the 14 C57BL/6 and C57BL/10 mouse lines. On average, each line had an estimated 1.27 Mb of simple repeats in total (range 1.07 – 1.47 Mb), or 0.051% of the genome. In the version 39 genome assembly, we found 25 Mb (1% of the genome) of simple repeats meeting the same parameters as could be found by k-Seek with short reads. This likely indicates biases against sequencing simple tandem repeats with Illumina, which we found previously (Flynn et al. 2020). Most simple tandem repeats in the assembly were contained in arrays less than 1kb long. However, we believe that our sample of simple tandem repeats found with short read sequencing is still informative and likely representative of the whole genome.

There were 434 kmers that had at least 10 copies in at least one line (Table S1). 63 kmers had an average abundance of at least 100 copies across the lines (Table S2), which we refer to as “common kmers.” There was no obvious clustering of samples by relatedness on a PCA of kmer abundances on PC1 or PC2, but there was some clustering of the B6 and B10 clades on PC3 (Figure S2). The most common kmer unit length was 6mers, with 22/63 common kmers being 6mers. 4-and 5-mers are also common, composing 11 and 8 of the common kmers, respectively. The GC content of kmers has a median of 0.5 and an average of 0.44, slightly higher than the genome-wide average GC content of 0.41. As in other species, several repeats were related to others by inferred single or few mutational steps (Figure 1). This fits the model of a novel repeat sequence being seeded by a mutation in an existing repeat, and then expanding into its own array. Extensive derivation of similar kmers (most 5 or 6 bp) is especially common for AC-only and AG-only repeats (Fig 1). AC and AG-only repeats make up a total of 25% of the total simple repeat abundance (as measured by total number of base pairs) in the mouse genome.

**Figure 1:**
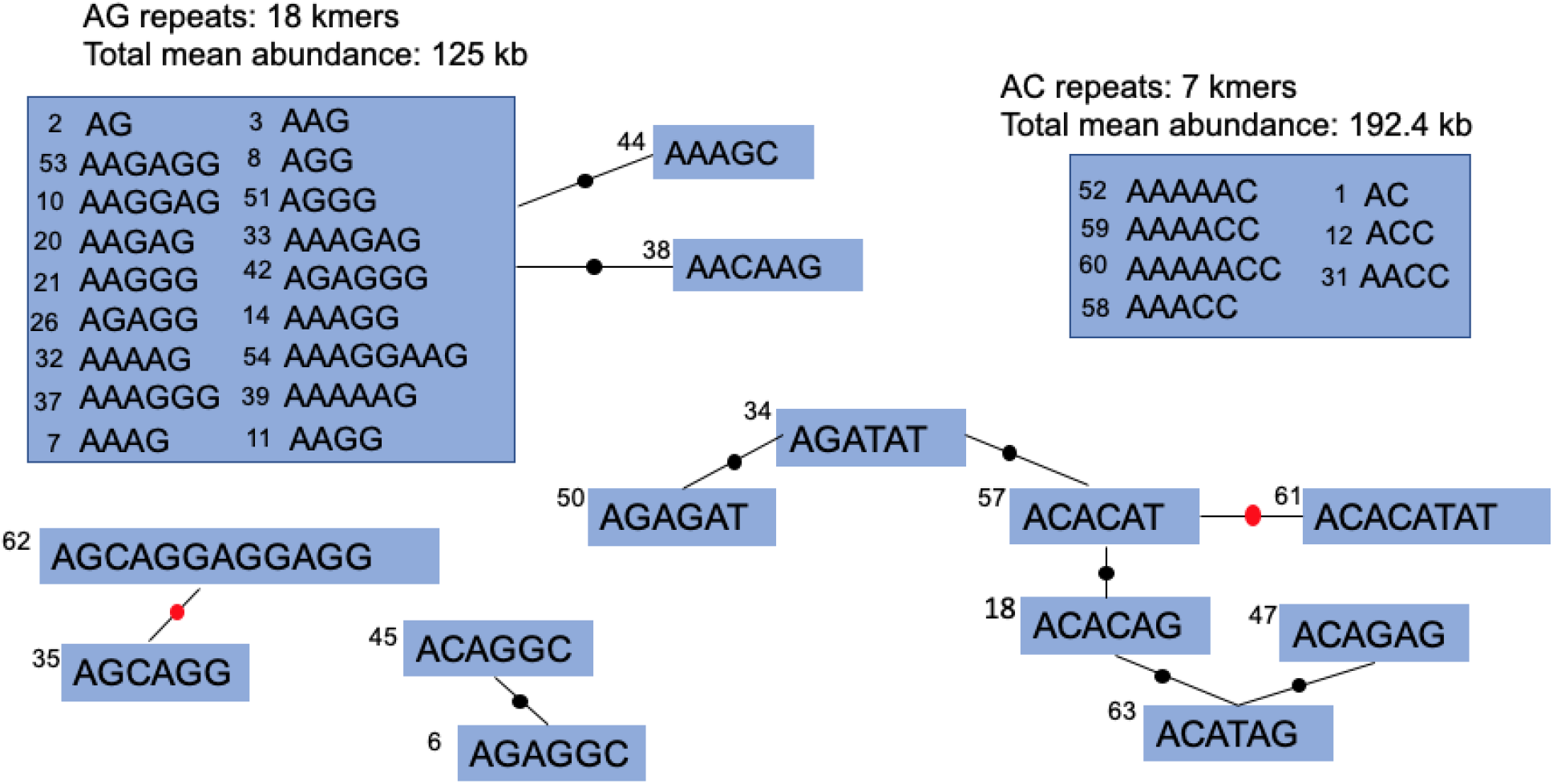
Relatedness diagram of abundant kmers in the C57BL/6 and C57BL/10 strains. AG and AC repeats are grouped together because there are many possible mutational steps that could derive these related sets of kmers. A black dot is a single nucleotide difference between the kmers, and a red dot is a copy number change or indel. Each kmer is labelled with a number, which represents its relative abundance (1 being the highest).

### Simple repeats vary in their rate of copy number changes

We calculated the difference in copy number from the inferred ancestral state at the nearest node and the terminal branch, and then divided by the number of generations between the node and the terminal branch (Figure 2A). The rate of copy number change can be positive or negative depending on whether the line experienced a net gain or loss of repeat copy number since the ancestor. Copy number change rates varied in magnitude and directionality among kmers (Figure 2B). However, across all kmers there was not an overall directional trend (Figure 2B), in contrast to the pattern we found in Daphnia, where most kmers tended to increase in abundance (Flynn *et al*. 2017). The overall abundance of kmers varied across the phylogeny, but did not follow the tree topology (Figure 2C).

**Figure 2:**
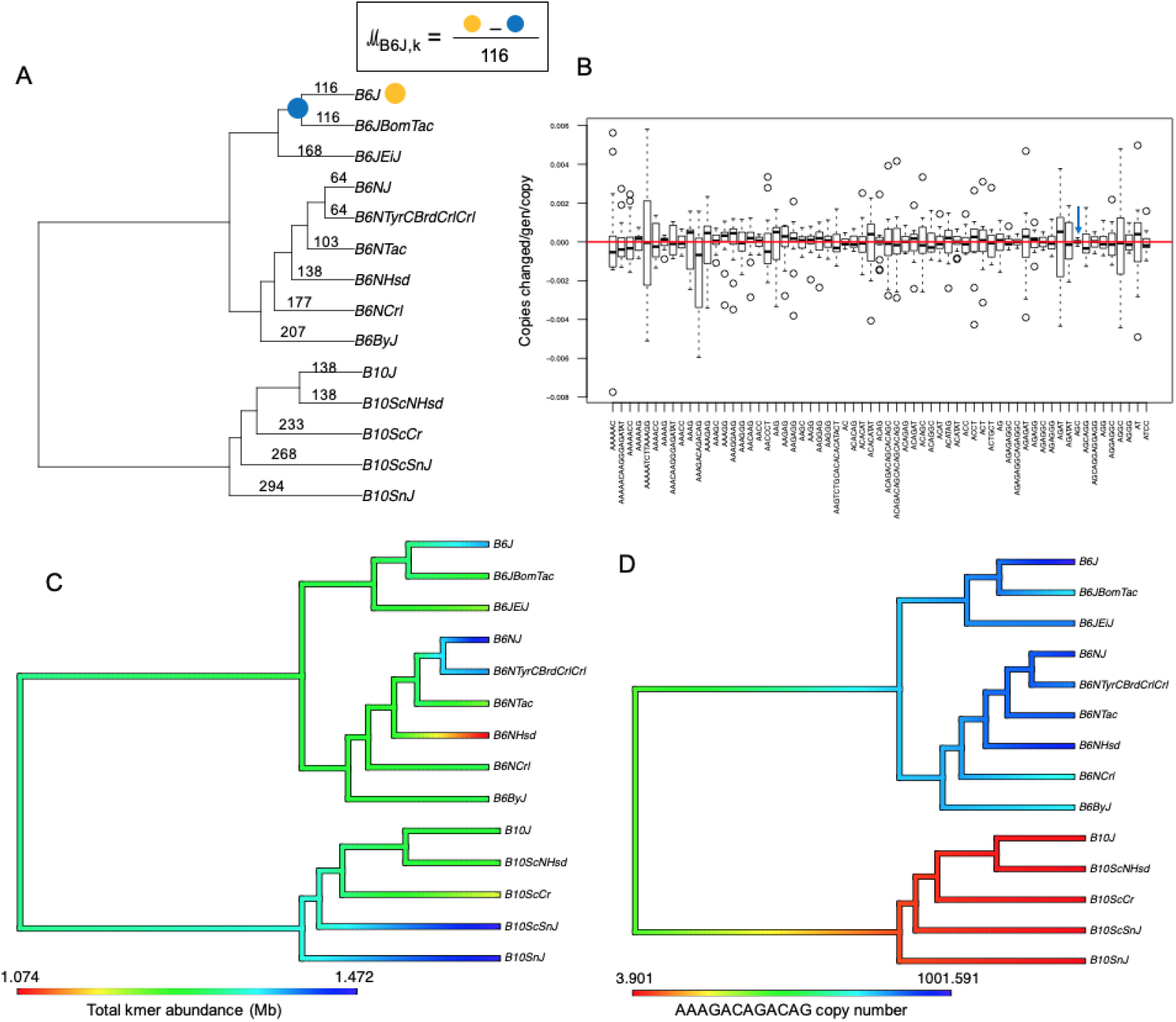
Copy number change rates in mouse simple repeats. Line names are abbreviated to reduce redundancy and space. A) The phylogeny based on historical records, with calculation of mutation rates being illustrated for the C57BL/6J line. B) Box plot of per kmer normalized mutation rate across all phylogenetic branches. The red line passes through zero, indicating an overall balance in the directionality of mutation rate. The blue arrow highlights AGC (or CAG) repeats, which have the lowest mutation rate of all kmers. C) Phytools plot showing variation in total kmer abundance within each line. Color gradients are based on the assumption of Brownian motion by Phytools package. D) An example of a kmer displaying a strong phylogenetic signal.

Of the 64 common kmers, AGC has the lowest absolute (either positive or negative) copy number change rate, only varying between 572 and 613 copies among all 14 lines (Figure 2B). AGC (or CAG) repeats are sometimes found in long arrays in protein-coding genes and cause several Mendelian disorders (Lieberman *et al*. 2019). Given the extremely low variance in copy number among lines despite hundreds of generations, we hypothesize selection has influenced the maintenance of copy number. Mice with phenotypic defects, such as those caused by CAG expansions in coding regions, would not be used to propagate the line. AAAAATCTTAAAGG had the highest mutation rate of all kmers with an absolute normalized copy number change rate of 2.2 x 10^−3^ copy changes per generation per copy. The abundance of the telomere repeat (AACCCT) is expected to vary from within-generation telomere shortening and telomerase activity, rather than solely inter-generational copy number changes. This repeat ranged in abundance among lines by a factor of two, from 22,781 to 46,030 copies, but we cannot rule out that this variation is due to imperfect age-matching across samples.

Most kmer abundances were not more similar between lines that are closely related than distantly related. However, we found several kmers that had a phylogenetic signal (Table 1, Figure 2D). Almost all cases of phylogenetic signal showed grouping/separation in copy number between the two major clades C57BL/6 and C57BL/10. This may be driven by the detected introgression from a divergent strain of about 5% of the genome likely into C57BL/10 (Mortazavi *et al*. 2020). Of the 7 kmers with a strong phylogenetic signal, 3 represented losses in C57BL/10 and gains in C57BL/6, and 3 represented gains in C57BL/10 and losses in C57BL/6, and one was a gain specific to the C57B/6J clade.

**Table 1:** kmers displaying a phylogenetic signal of abundance in the tree.

| Kmer | C57BL/6 pattern | C57BL/10 pattern |
| --- | --- | --- |
| AAAGACAGACAG | 700-1000 copies | <5 copies |
| AAAGACAG | 50-95 copies | 0-2 copies |
| AGCATGG | < 4 copies | 10-17 copies |
| AAAGAAAGG | 50-200 copies (1 outlier 10 copies) | 4-27 copies |
| AGGGC | C57B/6J clade 51-57 copies, B6N/By clade 56-88 copies | 84-190 copies |
| AAGAAGAAGAGG | 35-70 copies | 64-230 copies |
| AAAGAAAGGAAGGAAGG | C57BL/6J clade 13-18 copies, C57BL/6N-By clade 0-10 copies | 2-10 copies |

### Inbred lines varied greatly in their rates of genome-wide simple repeat copy number changes

Copy number change rates also varied in magnitude and overall direction on a per line basis. Although the magnitude of rate of change tended to be correlated with copy number, the slope and direction of change varied across lines (Figure 3). Nine of the 14 lines had a significant deviation from 50% of kmers having a net increase and 50% having a net decrease by the sign test, which would be expected if the gains and losses of each kmer were equally likely and independent from each other. In particular, some lines had consistent patterns of either gain or loss of most simple repeats, and these patterns ranged from relatively modest trends to high magnitude patterns. For example, in C57BL/6NCrl, the kmers with higher abundance have higher copy number change rates, but these were roughly symmetrical in positive and negative directions (Figure 3A). In C57BL/10ScSnJ, almost all kmers had positive rates, but the magnitude of change was low (Figure 3B). C57BL/6NJ experienced a mostly consistent positive rate with several gains at high magnitudes (Figure 3C), corresponding to a high positive change rate of +8.1 x 10^−4^ copies gained per copy per generation. C57BL/6NHsd, experienced both a consistent loss and high magnitude (Figure 3D), corresponding to a high negative copy change rate on average across all kmers of −9.11 x 10^−4^ copies lost per copy per generation. Since their separation an estimated 138 generations ago, C57BL/6NHsd lost 190 kb (16%) and C57BL/6NJ gained 205 kb (17.7%) of total genomic simple repeats compared to their common ancestor. These lines are potentially simple tandem repeat hypermutators, similar to what we found in Chlamydomonas (Flynn et al. 2018). Known single-nucleotide hypermutators in mice have defects in repair machinery (Uchimera et al. 2015), and hypermutators are rarely found spontaneously in the wild or laboratory populations in mammals. C57BL/6NJ and C57BL/6NHsd are especially intriguing because the cause of higher repeat copy change rate is unknown, and there is a consistent directionality of either simple repeat losses or gains. Different repair/monitoring mechanisms may be involved in expansions versus contractions, or variants in heterochromatin genes or other heterochromatic elements such as complex satellites or TEs may result in either losses or gains being favored.

**Figure 3.**
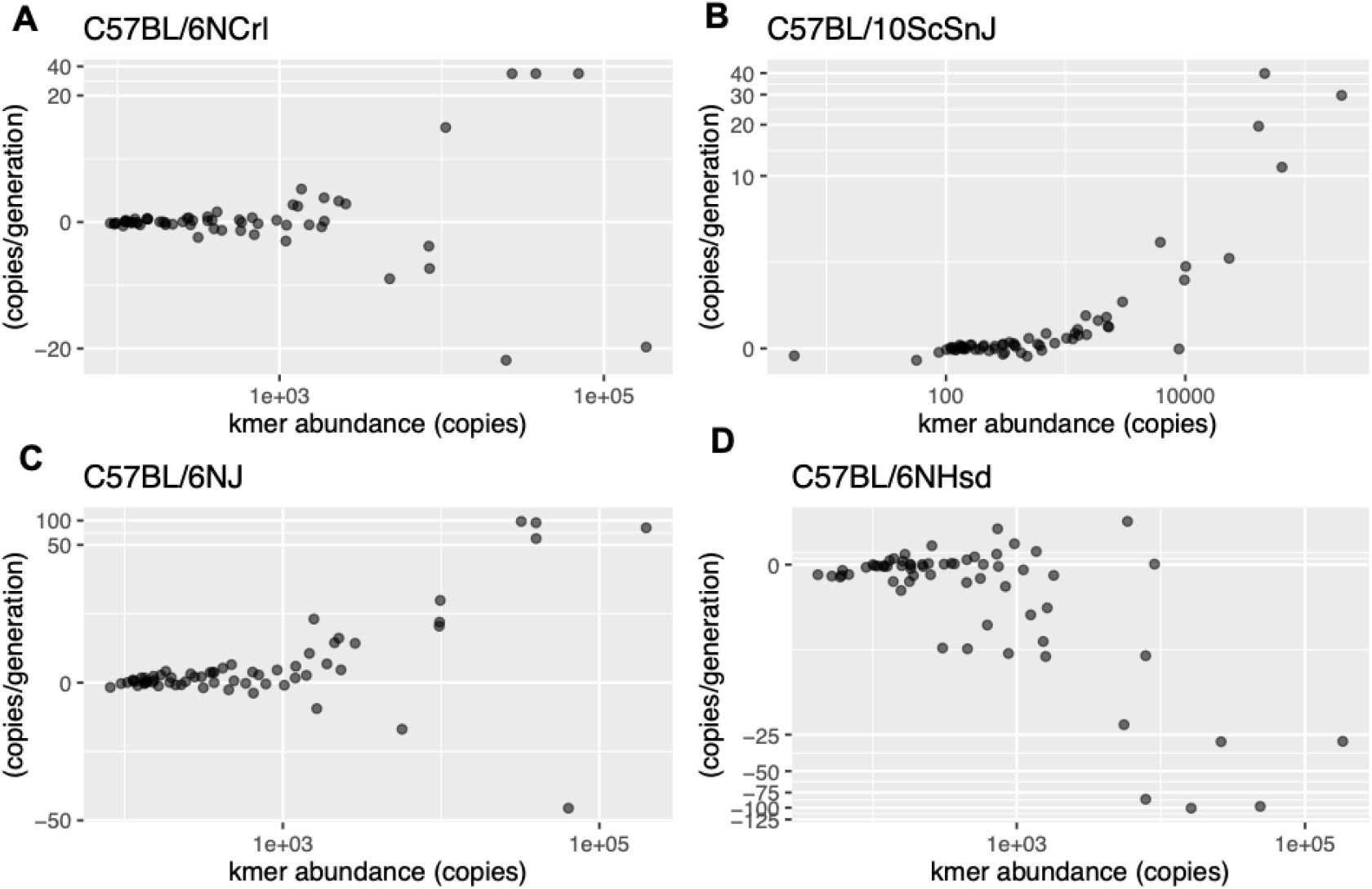
Relationship between kmer abundance and copy number change rate in four different inbred mouse strains. Each dot is a distinct kmer repeat. The X axis is on a log10 scale and the Y axis is on a pseudo-log10 scale, to allow scaling of negative rates, using the R package ggalin.

We also calculated absolute copy number change rates, where we took the absolute value of the directional rate. The overall average absolute rate across all lines and kmers was 7.28 x 10^−4^, slightly lower than that of Daphnia at 2.74 x 10^−3^ (Flynn *et al*. 2017) and slightly higher than that of Chlamydomonas (Flynn *et al*. 2018).

### Complex pericentromeric satellites have similar mutation rates to simple repeats

The mouse genome contains two abundant complex satellites called the major and the minor satellite. The major satellite is located in the pericentromeric region of all mouse chromosomes except the Y and has unit length 234 bp, while the minor satellite is located at the centromeres of all mouse chromosomes (except the Y) and has unit length 120 bp (Guenatri et al. 2004). We also estimated the copy number change rates of the major (Figure 4B) and minor satellite based on estimated copy number from average read mapping depth. The major satellite’s estimated copy number ranged from 923,277 – 1,317,473 among the 14 lines, while the minor satellite’s estimated copy number ranged from 123,362 - 143,527. The major satellite copy number change rate was 4.8 x 10^−4^ copies/generation/copy, slightly lower than that of simple repeats. The minor satellite change rate was less than half that of the major at 1.9 x 10^−4^ copies/generation/copy. Interestingly, these rates are within 2-fold of the simple repeat rates.

**Figure 4.**
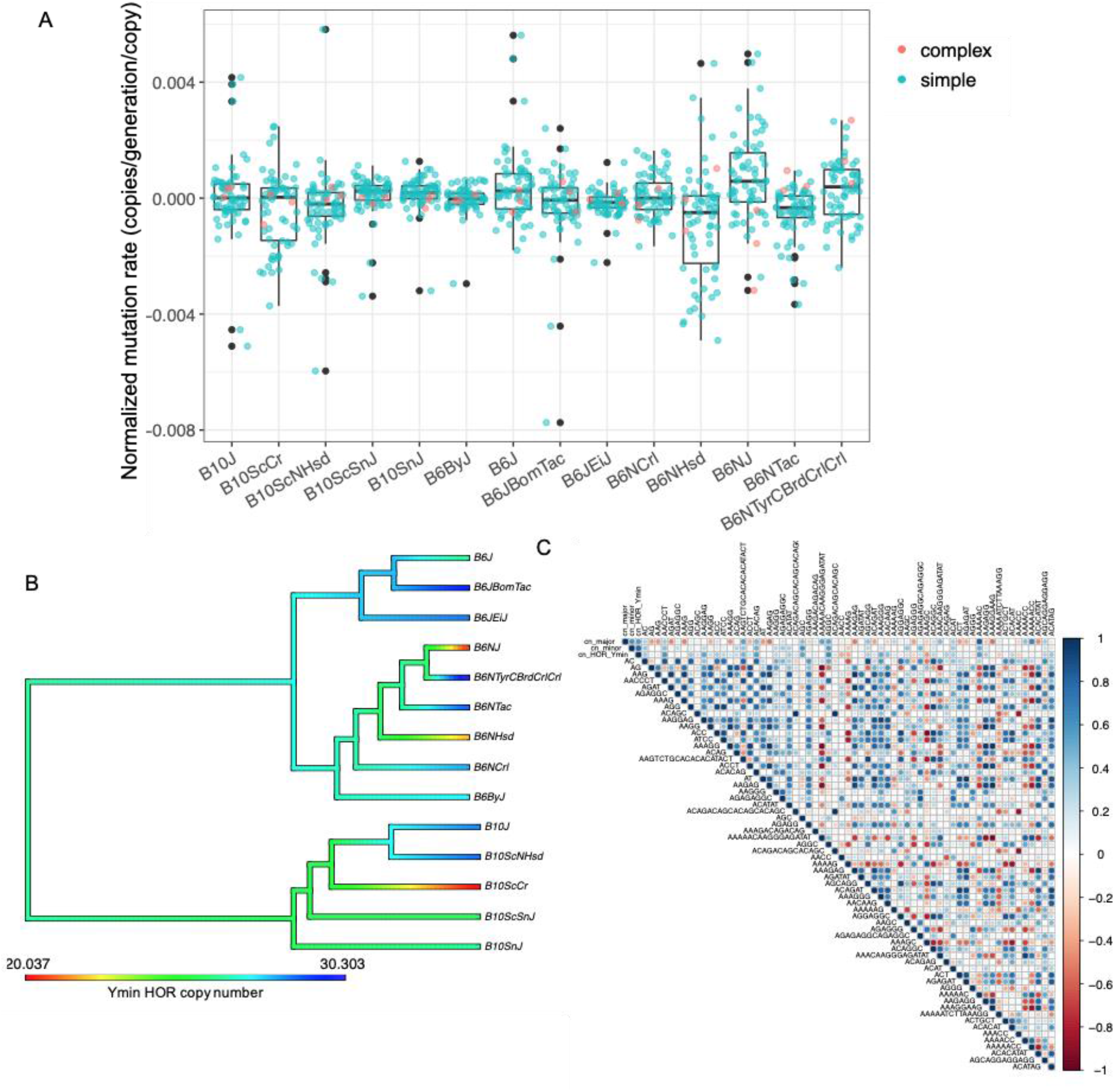
Interplay of simple and complex tandem repeat mutation. Line names are abbreviated to reduce redundancy and space. A) Boxplot of per line mutation rate. The mutation rates are shown for each line, with simple repeats in blue and complex in orange. For line C57BL/6NHsd, there is a trend of overall negative simple repeat mutation rate while this is not the case for complex satellites. For line C57BL/6NJ, there is an overall positive mutation rate for simple repeats, but overall negative for complex satellites. B) Phytools plot showing variation in Ymin copy number across all lines. C) Correlation plot showing correlations between simple and complex repeat mutation rates across all lines.

The Y chromosome of mouse contains a highly diverged Y-specific minor satellite sequence, called Ymin. Ymin is arranged in higher order repeats (HORs), wherein each monomer of approximately 121 bases within the HOR is highly diverged, but different HOR copies are very similar (Pertile *et al*. 2009). This structure is similar to the human alpha satellite (Sullivan *et al*. 2017), but does not occur on the other chromosomes of the mouse genome. A previous study showed with pulsed-field gel electrophoresis (PFGE) that no major changes in Ymin structure have occurred among inbred lines, including B6, over hundreds of generations. However, PFGE has much lower sensitivity than whole-genome sequencing, which we underscore here. We estimated HOR copy numbers of Ymin in all our sequenced strains. We found that the mutation rate was variable among lines. Interestingly, two closely related lines had highly diverged copy numbers that resulted in mutation rate estimates of +/- 110 bp per generation, which amounts to on average almost a full monomer unit lost every generation. Overall, the per copy number (HOR) change rate of Ymin was 4x higher than the per copy number (monomer) change rate of the non-Y centromeric minor satellite. Previous work has found that high mutation rates on human Y chromosomes drive high levels of structural variation (Repping *et al*. 2006). Furthermore, Ymin has a complex HOR structure whereas the minor centromeric satellite is composed of more homogeneous monomers, and based on these structures may have different constraints in copy number change mechanisms. Ymin, only present on a single haploid chromosome, is hypothesized to evolve exclusively by out-of-register sister chromatid exchange (Pertile *et al*. 2009), whereas the minor satellite likely also evolves by homologous and non-homologous recombination, driving homogenization on all chromosomes except the Y (Kalitsis *et al*. 2006).

Interestingly, the lines that had extreme losses and gains in simple repeats experienced the opposite trend for complex satellites (Figure 4A). C57BL/6NHsd had the highest rate of loss of simple repeats and had the largest gain in the major and minor satellites (major gain: 129958 copies, minor loss: 666 copies). C57BL/6NJ, which had the highest rate of gain in simple repeats had the largest loss in major and minor satellites (major loss: 126611, minor loss: 1253) (Figure 4A). This observation indicates that hypermutators in simple repeats are not hypermutators in complex repeats, suggesting that the dominant molecular machinery mediating these events are distinct. This is concordant with findings and assumptions in the field. Simple repeats are often assumed to be dominated by replication slippage (Li *et al*. 2002), while complex repeats are likely to evolve by recombination (Henikoff 2002; Kalitsis *et al*. 2006). Additionally, the different patterns in simple versus complex repeats may reflect compensation under a model of stabilizing selection on genome-wide repeat abundance (Flynn *et al*. 2017). These two lines also did not have higher rates of SNPs or small indels than other lines (Mortazavi et al. 2020). However, C57BL/6NHsd had the highest number of large-scale deletions found by CNVnator, and C57BL/6NJ had the highest number of duplications called by Lumpy (Mortazavi et al. 2020), concordant with the direction of gains and losses of kmers that we found.

We constructed a correlation matrix of directional mutation rates including the top 63 simple tandem repeats and the three main complex satellites (Figure 4C). Positive correlations can indicate kmers located physically close to each other, and negative correlations can indicate trade-offs or conflicts between different kmers. The complex satellites had little overall positive or negative correlations with any simple repeats or each other, despite apparent negative correlations in a small subset of lines (discussed above). Importantly, Ymin had no positive correlation with the minor satellite, indicating that the mechanisms through which each are changing in copy number are most likely independent. The repeats AAAAACAAGGGAGATAT, AAAAACC, AAAACC, and AAAAG, had strong negative correlations with many other kmers in the genome. These comprehensive negative correlations were not common in Daphnia and Chlamydomonas and would require further study to understand the mechanistic causes behind these patterns. These kmers may be evolving under net copy number constraints (possibly affecting a large portion of the genome) which manifest as negative correlations with other kmers.

## Conclusions

After analyzing simple and complex tandem repeat copy numbers in 14 inbred mouse strains, we discovered a variety of interesting patterns. AC and AG kmers dominate the simple repetitive portion of the genome, and like in other species, there are inter-connected networks of relatedness among simple repeat motifs. Copy number change rates of simple repeats varied across both individual strains and repeat motifs. The AGC (or CAG) repeat had the lowest net rate of copy change, with almost identical copy numbers among lines, suggesting this repeat (which is often found in coding regions), may be under stabilizing selection. There were two strains that had particularly high rates of change that were in consistent but opposite directions (gain and loss) across genome-wide simple repeats. Those high and directional rates were not applicable to the major and minor satellites, suggesting that different mutational mechanisms are dominant for simple versus complex repeats in mouse. Once normalized by copy number, we found that the copy number rates of change were very similar for these two types of tandem repeats (within 2-fold). As a comparison, other mutation types like single nucleotide substitutions and indels, vary by five orders of magnitude in mouse (Kruglyak *et al*. 1998; Uchimura *et al*. 2015). This consistency of rate of change may reflect inherent properties of the mutational processes or genomic constraints. The Y chromosome in mouse contains a diverged satellite called Ymin that contains higher order repeats, a structure unlike the centromeric repeats on the other chromosomes (Pertile *et al*. 2009). Ymin had a four-fold higher mutation rate than the homologous minor satellite on all chromosomes except the Y, despite being only on a single haploid chromosome and having more limited types of recombination that can occur. Finally, some simple repeats had strong negative correlations in their copy number change rates with many other repeats genome-wide, likely implicating yet to be understood constraints.

## Data availability

Raw reads are available from NCBI SRA under accession PRJNA705216. Scripts and input data required to reproduce our results are available at: https://github.com/jmf422/BL6_mouse_satellites

## Acknowledgements

We would like to thank Elissa Cosgrove for help with reproducing the phylogenetic tree. We would also like to thank Abraham Palmer, Yangsu Ren, Milad Mortazavi, and Melissa Gymrek for helpful discussions. Ian Caldas gave advice on implementing the GC correction scripts. This work was supported by NIH R01 GM119125 to Andrew G. Clark and Daniel Barbash.

